# Genotype-matched Newcastle disease virus vaccine confers improved protection against genotype XII challenge: The importance of cytoplasmic tails in viral replication and vaccine design

**DOI:** 10.1101/492421

**Authors:** Ray Izquierdo-Lara, Ana Chumbe, Katherine Calderón, Manolo Fernández-Díaz, Vikram N. Vakharia

## Abstract

Although typical Newcastle disease virus (NDV) vaccines can prevent mortality, they are not effective preventing viral shedding. To overcome this, genotype-matched vaccines have been proposed. To date, this approach has never been tested against genotype XII strains. In this study, we generated and assessed the protection against genotype XII challenge of two chimeric NDV vaccine strains (rLS1-XII-1 and rLS1-XII-2). The rLS1-XII-1 virus has the complete fusion protein (F) and the hemmaglutinin-neuraminidase (HN) open reading frames replaced with those from genotype XII strain NDV/peacock/Peru/2011 (PP2011) in a recombinant LaSota (rLS1) backbone. For rLS1-XII-2 cytoplasmic tails of F and HN proteins were restored to those of rLS1. *In vitro* studies showed that rLS1-XII-2 and the parental rLS1 strains replicate at higher efficiencies than rLS1-XII-1. In the first vaccine/challenge experiment, SPF chickens vaccinated with rLS1-XII-1 virus showed only 71.3% protection, whereas, rLS1 and rLS1-XII-2 vaccinated chickens were fully protected. In a second experiment, both rLS1-XII-2 and the commercial vaccine strain LaSota induced 100% protection. However, rLS1-XII-2 virus significantly reduced viral shedding, both in the number of shedding birds and in quantity of shed virus. In conclusion, we have developed a vaccine candidate capable of fully protecting chickens against genotype XII challenges. Furthermore, we have shown the importance of cytoplasmic tails in virus replication and vaccine competence.

## Introduction

Newcastle disease virus (NDV) is a widely distributed virus that affects poultry and other avian species [1]. NDV belongs to the order of Mononegavirals, family Paramyxoviridae and genus Avulavirus [2]. NDV, formerly known as the Avian Paramyxovirus type-1, is formally known as the *Avian Avulavirus 1* (AAvV-1) since 2016 [https://talk.ictvonline.org/taxonomy/]. NDV has a nonsegmented single-stranded negative-sense RNA genome of 15,186 bp in length, which follows the rule-of-six [3]. NDV genome encodes six structural genes: Nucleoprotein (N), phosphoprotein (P), matrix (M), fusion (F), hemagglutinin-neuraminidase (HN) and large polymerase (L) [4]. From these proteins, M, HN and F form the envelope. The M protein is located at the inner face of the viral membrane and is responsible to drive the viral budding and virion assembly process [5]. HN and F proteins are surface glycoproteins anchored to the viral envelope. Both HN and F are incorporated into the virions via the interaction of their cytoplasmic tails with the M protein [6,7]. The HN protein mediates the attachment of the virus to the host cell receptor, and the F protein mediates fusion of viral and host cell membranes [3]. The F protein requires to be cleaved into F1 and F2 prior to fusion with cell membranes [8]. The F protein cleavage site of avirulent (lentogenic) strains exhibit a dibasic motif (i.e. ^112^GRQGRL^117^), while virulent (mesogenic and velogenic) strains exhibit a polybasic motif (i.e. ^112^RRQKRF^117^) [8,9].

Based on the complete sequence of the F gene, NDV strains are classified into two classes I and II [10]. Class I contains a single genotype, and strains have been isolated mainly from wild birds and are generally lentogenic [10]. Class II contains at least 18 genotypes (I-XVIII), and they can be lentogenic, mesogenic or velogenic [10,11]. Based on Diel et al. (2012) classification rules, an evolutionary distance between 3% and 10% among clades within a genotype allows its subdivision into subgenotypes [10]. Commonly vaccine strains (i.e. LaSota) belong to genotypes I and II and are used all over the world. On the other hand, genotype XII strains are highly virulent and have been isolated from Peru, Colombia, China and Vietnam [12–15]. So far, at least three subgenotypes are distinguished within genotype XII: XIIa, XIIb and XIId. Subgenotype XIIa strains have been isolated only in South America (Peru and Colombia) [13–15], XIIb strains have been isolated only in the province of Guangdong in China [10,16] and XIId strains were recently reported in Vietnam [12]. XIIc is a potential subgenotype composed of strains isolated between 1986 and 2005, nevertheless only partial sequences of the F gene are available for these strains, therefore it cannot be consider as a proper subgenotype yet [13]. In Peru, XIIa is the only genotype isolated so far [13] and despite intensive vaccination campaigns, several outbreaks are reported every year [17], even in vaccinated flocks. This can be explained by antigenic differences between vaccine and genotype XII circulating strains. Amino acid identities of F and HN proteins within subgenotype XIIa is above 99%, and within overall genotype XII strains are above 90% (Table 1). While F and HN protein sequence identities between XIIa and vaccine strains, as LaSota, are only between 80 and 90% (Table 1). Hence, it is reasonable to think that vaccines could be improved by matching their antigenicity with those of field strains.

**Table 1.**
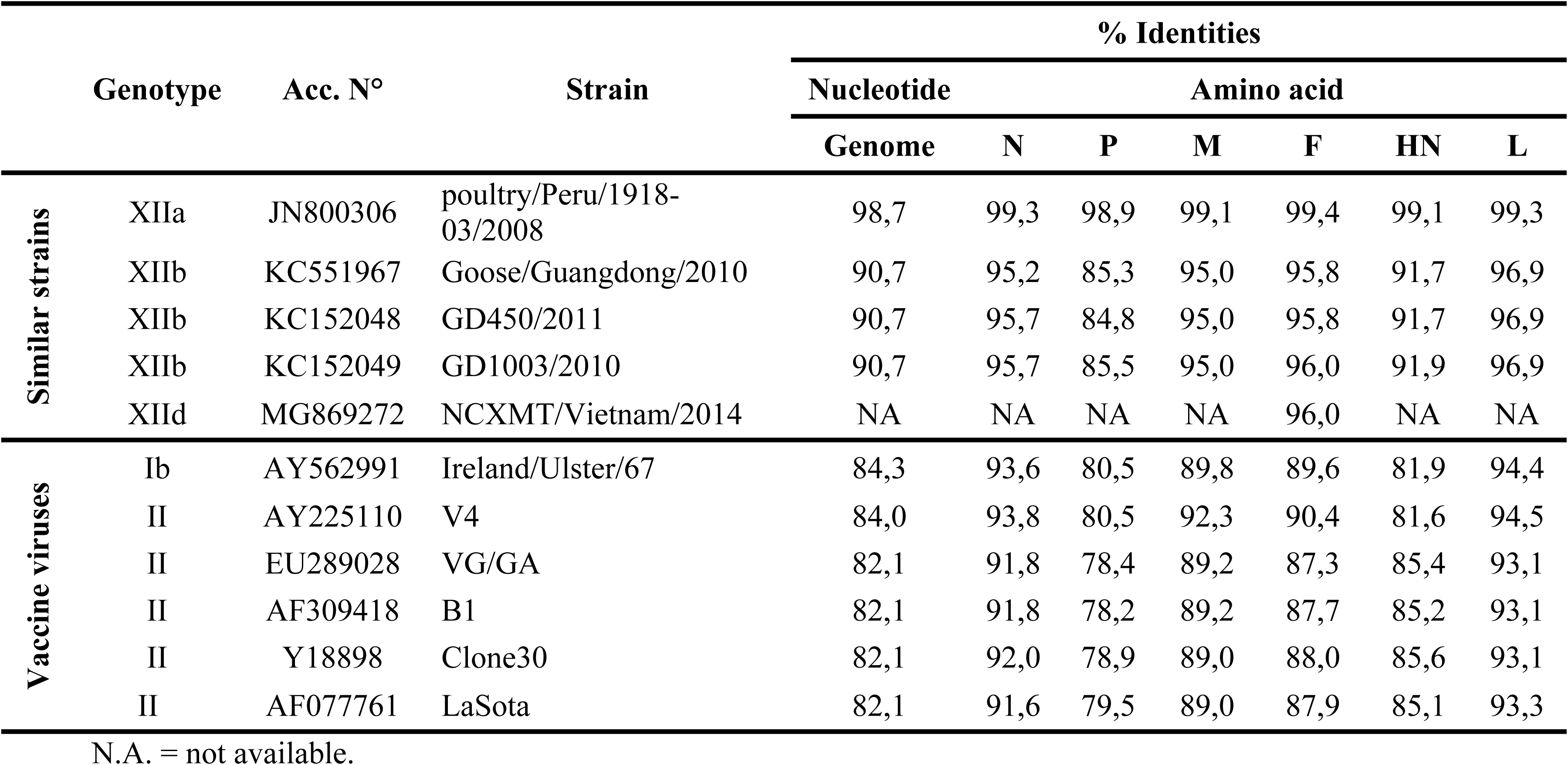
Sequence identities between PP2011 and other representative NDV strains.

In non-vaccinated chickens, virulent strains are capable to produce up to 100% morbidity and mortality [1]. While classical vaccines are capable to prevent the disease under experimental conditions, they fail to prevent viral shedding [18–20]. Many authors have suggested that antigenic matches of F and HN proteins between vaccine and challenge strains are capable to improve vaccine protection by significantly reducing viral sheading [21,22]. However, some other authors have suggested that genotype mismatch is not the main reason for vaccination failure in immunocompetent chickens [23,24]. To date, live-attenuated genotype-matched vaccines have been tested for genotypes V, VII and XI [21]– [27]. However, no data has been recorded whether genotype XII homologous vaccine is capable to induce a better protection than commonly used vaccines.

To assess whether genotype XII matched vaccine can improve protection against a homologous challenge, we generated by reverse genetics two recombinant NDV (rNDV) strains containing both F and HN proteins from the genotype XIIa strain NDV/peacock/Peru/2011 (PP2011) [14] in the rLS1 backbone [28]. In the first rNDV, rLS1-XII-1 virus, complete F and HN open reading frames (ORFs) were replaced with those from the PP2011. For the second virus, rLS1-XII-2, only the ecto- and transmembrane domains of the F and HN proteins were replaced in rLS1. To reduce pathogenicity in both viruses, the F cleavage site was changed from the polybasic motif ^112^RRQKRF^117^ to the dibasic motif ^112^GRQGRL^117^. In the first vaccine/challenge experiment, rLS1-XII-1 and -2 viruses were compared against the parental rLS1. In the second experiment, rLS1-XII-2 virus and the commercial vaccine LaSota strain were compared for its capacity to induce protection, specific antibodies and reduce viral shedding.

## Materials and methods

### Cell lines, viruses and animals

DF-1 (derived from chicken fibroblasts) were maintained in Dulbecco’s modified Eagle medium (DMEM) F12 (HyClone), supplemented with 5% heat-inactivated fetal bovine serum (FBS) and 2.5% chicken serum (ChkS) (Sigma–Aldrich). Vero (monkey kidney) cells were maintained in DMEM F12 supplemented with 5% FBS. Both cells lines were cultured with 100 U/mL of penicillin and 100 µg/mL of streptomycin at 37 °C in an atmosphere of 5% CO2. The virulent strain NDV/peacock/Peru/2011 (PP2011) (Genbank accession number: KR732614), belonging to genotype XII, was previously isolated in Peru [14]. We previously developed the recombinant rLS1 vector [28]. The rLS1-XII-1 and rLS1-XII-2 were engineered and rescued in this study using the rLS1 strain as a backbone vector. All viruses were grown in 9- or 10- old specific pathogen free (SPF) chicken eggs (Charles River Laboratories, Wilmington, MA, USA). After 3 days of infection or egg death, allantoic fluids (AFs) were harvested, clarified, aliquoted, and stored at -80 °C. Titration was performed by median tissue culture infectious dose (TCID_50_). All protocols related to animal use were performed under the guidelines of the Committee of Ethics and Animal Welfare (CEBA) of the Faculty of Veterinary of the Universidad Nacional Mayor de San Marcos (UNMSM), Lima, Peru.

### Plasmid construction and virus recovery

The previously constructed pFLC-LS1 plasmid [28], containing the rLS1 genome, was used as backbone to clone rLS1-XII-1 and rLS1-XII-2 genomes. Fragments containing F and HN genes were chemically synthetized (Genscript, New Jersey, USA). The F_0_-XII fragment contains the complete ORF of the F protein from PP2011 strain and is flanked by the intergenic regions of rLS1 strain, including BssHII and MluI restriction sites at the ends. Five nucleotides in the cleavage site were changed to reduce pathogenicity, as consequence the polybasic motif ^112^RRQKRF^117^ was replaced by the dibasic motif ^112^GRQGRL^117^. The HN-XII fragment contains, from 5´ to 3´, a part of the F-HN intergenic region of the rLS1 parental vector, the complete ORF of the HN protein from PP2011 strain and the complete HN-L intergenic region, plus part of the L ORF of the rLS1 strain and is flanked by MluI and BsiWI restriction sites. Both F_0_-XII and HN-XII were sequentially inserted in pFLC-LS1 vector to generate the pLS1-XII-1 plasmid. F0-XII/CT-II and HN-XII/CT-II fragments were designed to contain similar characteristics as F0-XII and HN-XII, respectively, but with the cytoplasmic tails from the parental rLS1 vector (Fig 1). F_0_-XII/CT-II differs from F_0_-XII in that the sequence corresponding to the cytoplasmic tail of the F gene (29 amino acids at the C-terminal) was replaced by the one from the rLS1 strain. HN-XII/CT-II differs from HN-XII in that the sequence corresponding to the cytoplasmic tail of HN gene (22 amino acids at the N-terminal) was replaced by the one from the rLS1 strain. F_0_-XII/CT-II and HN-XII/CT-II fragments were sequentially cloned into pFLC-LS1 vector to generate the pLS1-XII-2 plasmid. All plasmids were purified using the Plasmid Mini Kit (QIAGEN), stored at -20 °C and sequenced. The rLS1-XII-1 and rLS1-XII-2 viruses were recovered as previously described [28]. Briefly, Vero cells plated at 90% confluence in 12-well plates were transfected with the supporting plasmids (pCI-N, pCI-P and pCI-L) plus either of the plasmids containing the viral genomes. On the next day, AF was added to a final concentration of 5% and cells were incubated for 4 days more. Cell supernatant was inoculated into 8-days old SPF chicken embryonated eggs and incubated for 4 days. AFs were harvested, clarified, and aliquoted and stored at -80 °C. Virus recovery was first confirmed by hemmaglutination assay (HA). To confirm the identity of the viruses, RNA was isolated from AF and a fragment of 708 bp was amplified and sequenced with primers F.1437_fw (5’-CAAGTTGGCAGAAAGCAACA-3’) and HN.296_rv (5’-GGATTCAAGTGCCACCTGTT-3’). Genetic maps of the parental virus (rLS1), the virulent wild-type NDV belonging to genotype XII (PP2011), and the genotype XII vaccine candidates (rLS1-XII-1 and rLS1-XII-2) are showed in Fig 1. The pathogenicity of the recovered viruses was determined by the intracerebral pathogenicity index (ICPI) and mean death time (MDT) were carried out on 1-day-old SPF chickens (Charles River Laboratories) and 10-day-old embryonated chicken eggs, respectively, using standard procedures [29].

**Fig 1.**
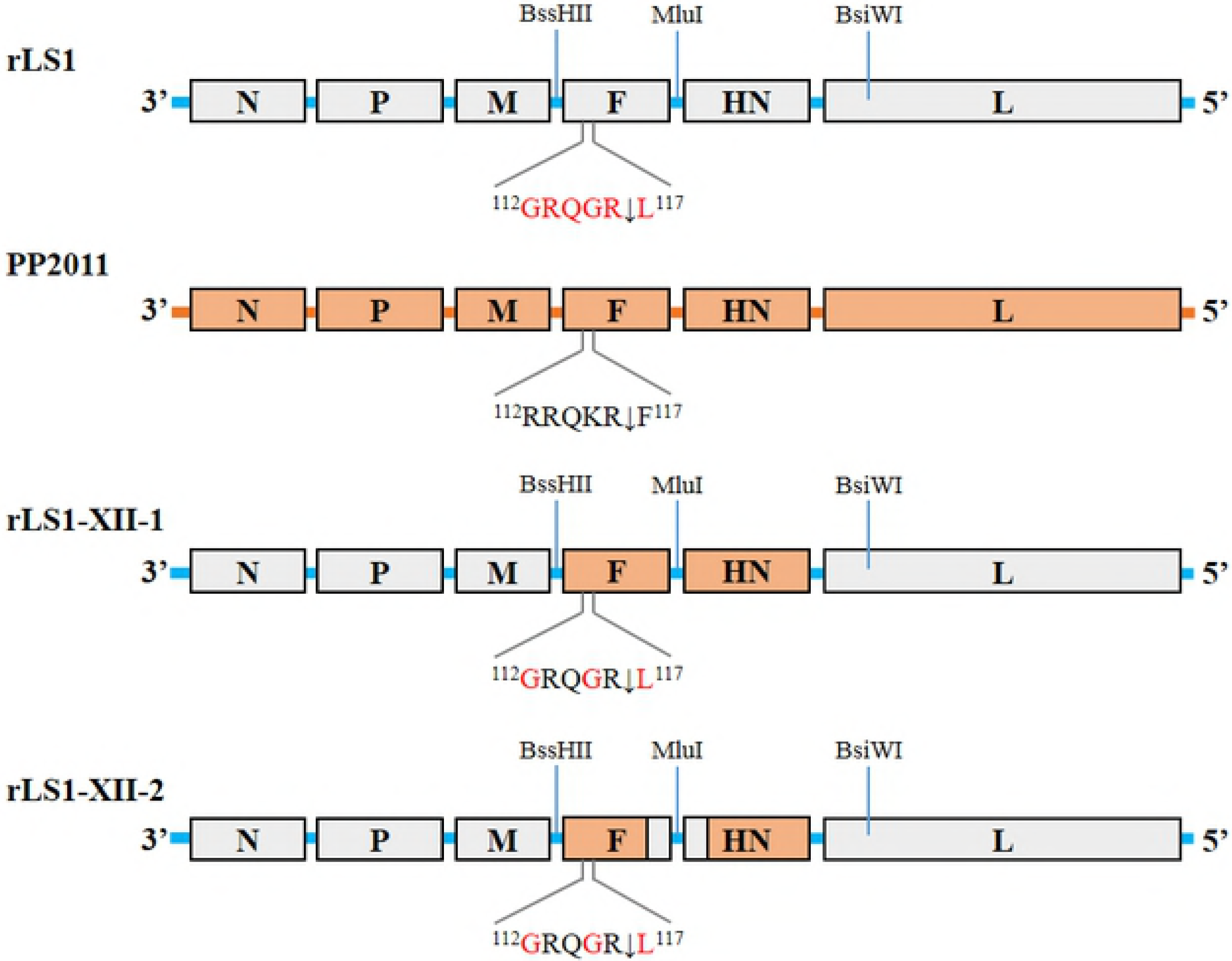
Viral gene maps. The gene map at the top is the parental NDV strain rLS1 and shows the three unique restriction sites used for swapping F and HN gene ORFs. The next map illustrates the wild-type strain PP2011 (genotype XII). The last two maps are the two chimeric derivative viruses. The rLS1-XII-1 has the complete F and HN ORFs swapped by those of the PP2011 strain, whereas rLS1-XII-2 has the cytoplasmic tails restored to those of the original rLS1 strain. The F protein cleavage sites are shown.

### *In vitro* growth kinetics in DF-1 cells

DF-1 cells were seeded at 50-60% confluence in 12-well plates and infected with rLS1, rLS1-XII-1 or rLS1-XII-2 at a multiplicity of infection (MOI) of 0.05. Cells were cultured with DMEM supplemented with 1% FBS and 5% AF with 5% CO2 at 37 °C. Supernatants were collected at 12, 24, 36, 48, 60 and 72 hours post-infection (hpi). Collected supernatants were quantified in DF-1 cells by TCID50. To clearly see the infection in monolayers, cells were stained with a monoclonal antibody against the NDV ribonucleoprotein (RNP) (cat. n° ab138719, Abcam, USA) as a first antibody and a goat polyclonal antibody anti-mouse IgG labeled with Alexa Fluor 594 (cat n° ab150116, Abcam) as a secondary antibody.

### Immunofluorescence

DF-1 cells were infected with either rLS1, rLS1-XII-1, rLS1-XII-2 viruses or mock-infected at a MOI of 0.01 for 1 hour, washed with Dulbecco’s phosphate-buffered saline (DPBS) and covered with DMEM semisolid media (0.75 % methylcellulose + 2 % FBS + 5% AF). Cells were incubated for 24, 48 and 72 hpi. Then, cells were washed 3 times with DPBS and fixed with 4% paraformaldehyde in DPBS for 15 min at room temperature (RT). Fixed cells were blocked with 5% Bovine Serum Albumin (BSA) (Sigma-Aldrich) in DPBS for 30 min at RT. Cells were incubated for 2 hours with a chicken polyclonal antibody against NDV (Charles River Laboratories) in a 1:2000 dilution in a solution of 5% BSA in DPBS. After washing, cells were incubated for 1 hour with goat anti-chicken IgY-Alexa Fluor^®^ 594 (cat n° ab ab150172, Abcam) at a final concentration of 1 µg/mL in a solution of 5% BSA in DPBS, and then washed. Cells were examined under the Observer.A1 fluorescence microscope (Carl Zeiss, Germany). Images were taken at 400X magnification with the AxioCam MRc5 camera (Carl Zeiss, Germany).

### First vaccination/challenge assay with recombinant vaccine viruses

A total of 52 one-day-old SPF chicks were divided into four groups. All groups were immunized twice at the 1st and the 14th days. Birds in the control group (n = 10) were mock-vaccinated with 30 µl of DPBS. Birds in groups rLS1 (n = 14), rLS1-XII-1 (n = 14) and rLS1-XII-2 (n = 14) were vaccinated via eye-drop route with 30 µl/bird of 10^7^ 50% egg infective dose (EID_50_/ml) of live rLS1, rLS1-XII-1 and rLS1-XII-2 viruses, respectively. At 28 days of age, chickens were challenged with 10^5^ median lethal dose (LD_50_) of PP2011 stock in 50 µl/bird. Chickens were observed for 10 days post-challenge (dpc). To assess viral shedding, oral and cloacal swabs were collected from all surviving chickens at 2, 4 and 7 dpc. Swabs were resuspended in 500 µl of DMEM media supplemented with 10% FBS and penicillin-streptomycin, then clarified by centrifugation at 20000 RCF per 10 min. 100 µl of each swap were inoculated into 10-day-old SPF chicken eggs and incubated for 3 days or until dead, then AFs were evaluated for HA to confirm presence of NDV.

### Second vaccine/challenge: Comparison between rLS1-XII-2 and the commercial LaSota vaccine strain

Based on the results of the first experiment, we decided to further continue with the evaluation of the rLS1-XII-2 and compare it against the commercial vaccine LaSota. A total of 38 seven-day-old SPF chickens were divided into three groups. Birds in the control group (n = 10) were mock-vaccinated with 30 µl of DPBS. Birds in groups LaSota (n = 14) and rLS1-XII-2 (n = 14) were vaccinated via eye-drop route with 30 µl/bird of live LaSota and rLS1-XII-2 (10^7^ EID_50_/ml), respectively. At 28 dpv, chickens were challenged with 10^5^ LD_50_ of PP2011 stock in 50 µl per bird. Chickens were observed for 10 days post-challenge (dpc). Serum samples were collected via wing-web bleeding at 2 and 4 weeks post vaccination. Antibody titers were calculated by neutralization test against either the challenge strain (PP2011) or a genotype II vaccine strain (rLS1-eGFP) as described previously [28]. Viral shedding was assessed in the same way as described above. Additionally, viral titers of HA positive swabs were calculated by plaque assay.

### Statistical Analysis

All statistical analysis were performed in GraphPad Prism 6.01 (GraphPad Software Inc., San Diego, CA).The differences in viral shedding between vaccinated groups were compared by t test. Mann-Whitney test was utilized to assess the differences in serum neutralization titers between vaccinated groups. Statistical significances were represented as *p < 0.05, **p < 0.01 and ***p < 0.001.

## Results

### Presence of homologous cytoplasmic tails in F and HN proteins results in a higher viral replication compared to heterologous cytoplasmic tails

In order to develop a genotype XII-matched vaccine, we engineered two chimeric NDVs. In rLS1-XII-1, complete F and HN genes were replaced in the rLS1 backbone for those of the PP2011 strain. In rLS1-XII-2 only the ecto- and transmembrane domains were replaced with those of PP2011. Both viruses were rescued and replicated in embryonated eggs at 10^8.8^ and 10^9.7^ EID_50_/ml, respectively (see Table 2). However, it is worth to mention that rLS1-XII-1 was detected by HA only after a second passage in embryonated eggs, while rLS1-XII-2 was HA-positive after only one passage (directly inoculated from transfection supernatant). Growth curves in DF-1 cells showed that parental rLS1 replicates 3.5 - 81.3 and 0 - 12.8 fold more compared to rLS1-XII-1 and rLS1-XII-2, respectively (Fig 2a). At the 36 hpi, there was no difference in the replication between rLS1 and rLS1-XII-2. Thus, in general, rLS1 replicates at higher titers than the rLS1-XII-2 and this one higher than rLS1-XII-1. Additionally, immunofluorescence evaluated at 24, 48 and 72 hpi showed that rLS1 and rLS1-XII-2 produce similar infection expansion patterns in monolayers of DF-1 cells, while rLS1-XII-1 have a much more restricted spread in cells (Fig 2b). Accordingly, no clear cytopathic effect (CPE) or syncytia formation was observed in monolayers infected with rLS1-XII-1, whereas rLS1 and rLS1-XII-2 presented the typical CPE of NDV. This can explain the differences in growth curves between these strains.

**Table 2.**
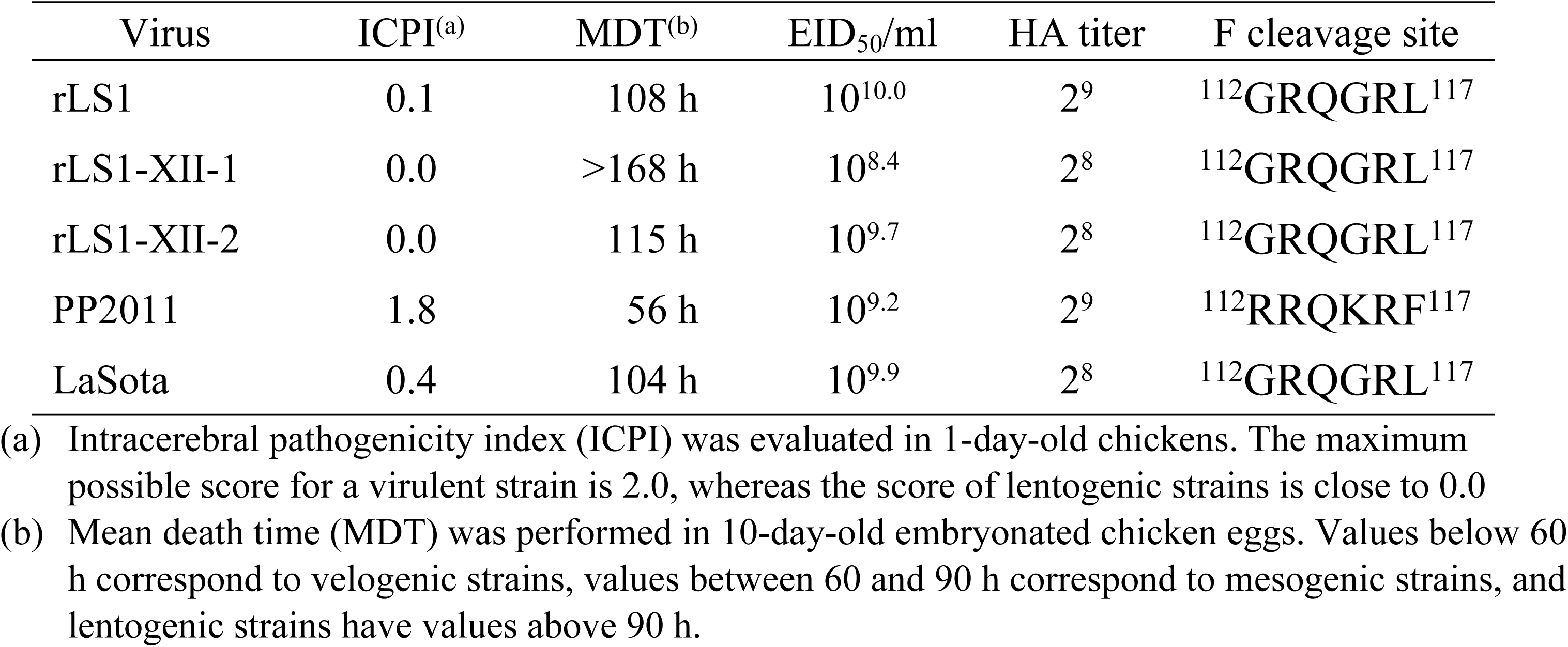
Biological characteristics of NDV strains used in this study.

| Virus | ICPI<sup>(a)</sup> | MDT<sup>(b)</sup> | EID<sub>50</sub>/ml | HA titer | F cleavage site |
| --- | --- | --- | --- | --- | --- |
| rLS1 | 0.1 | 108 h | 10<sup>10.0</sup> | 2<sup>9</sup> | <sup>112</sup>GRQGRL<sup>117</sup> |
| rLS1-XII-1 | 0.0 | >168 h | 10<sup>8.4</sup> | 2<sup>8</sup> | <sup>112</sup>GRQGRL<sup>117</sup> |
| rLS1-XII-2 | 0.0 | 115 h | 10<sup>9.7</sup> | 2<sup>8</sup> | <sup>112</sup>GRQGRL<sup>117</sup> |
| PP2011 | 1.8 | 56 h | 10<sup>9.2</sup> | 2<sup>9</sup> | <sup>112</sup>RRQKRF<sup>117</sup> |
| LaSota | 0.4 | 104 h | 10<sup>9.9</sup> | 2<sup>8</sup> | <sup>112</sup>GRQGRL<sup>117</sup> |

- (a) Intracerebral pathogenicity index (ICPI) was evaluated in 1-day-old chickens. The maximum possible score for a virulent strain is 2.0, whereas the score of lentogenic strains is close to 0.0 - (b) Mean death time (MDT) was performed in 10-day-old embryonated chicken eggs. Values below 60 h correspond to velogenic strains, values between 60 and 90 h correspond to mesogenic strains, and lentogenic strains have values above 90 h.

**Fig 2.**
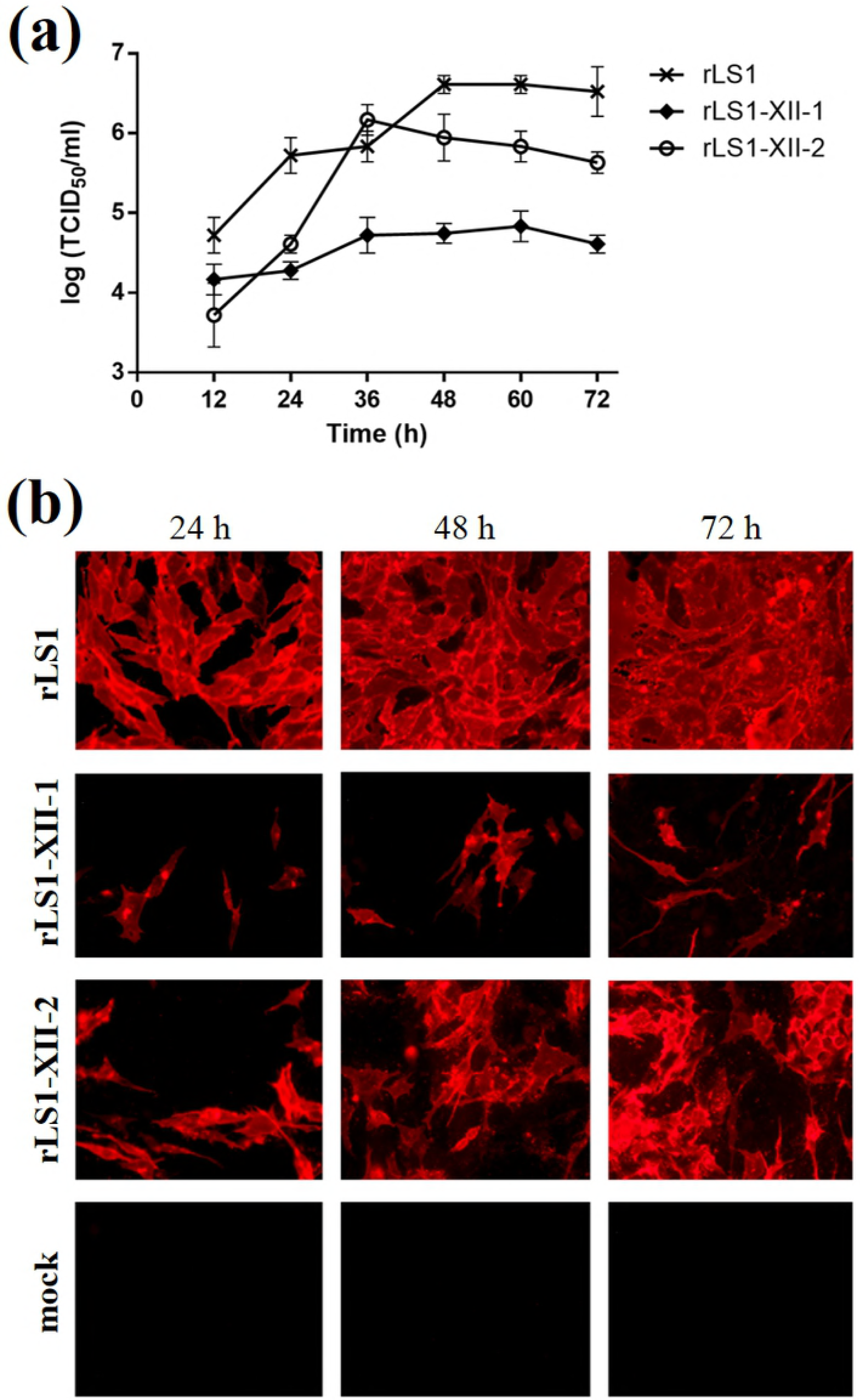
Characterization of recombinant viruses. (a) Multistep growth curves of rLS1, rLS1-XII-1 and rLS1-XII-2 in DF-1 cells. Cells were infected at a MOI of 0.05 and supernatants were collected in 12 hours intervals post-infection and tittered by TCID_50_/ml. Data was taken from three independent experiments. (b) In vitro infection patterns of the rNDVs. DF-1 cells were infected with a MOI of 0.01 for 1 hour and, then, covered with DMEM semisolid media containing 0.75 % methylcellulose. After 24, 48 and 72 h monolayers were washed and stained for NDV by immunofluorescence. All pictures were taken at 400X magnification.

The ICPI value for the parental rLS1 was 0.1, while for chimeric viruses rLS1-XII-1 and rLS1-XII-2 were 0.0 (Table 2). MDT values were in agreement with these results, where these three strains were classified as lentogenic (MDT > 90 hours). Although, rLS1-XII-1 showed the largest MDT (>168 h). PP2011 challenge strain was classified as velogenic due to its ICPI of 1.8 [14] and a MDT of 56 h. These results showed that replacing completely or only the transmembrane and ectodomains of the F and HN genes of rLS1 by those of the PP2011 did not increase the virulence of the vaccine strains, as long as a dibasic cleavage site is present in the F protein.

### The rLS1-XII-1 virus failed to protect chickens against a homologous challenge

In a first experiment, chickens immunized with two doses of rLS1, rLS1-XII-1 or rLS1-XII-2 were challenged at the four week of age with PP2011 strain (genotype XII) via oculo-nasal route. Both rLS1 and rLS1-XII-2 were capable of fully protecting chickens against mortality (14/14), but rLS1-XII-1 protected only 71.3 % (10/14), while mock-vaccinated control group did not protect birds against mortality (Fig 3). None of the chickens in the rLS1 or rLS1-XII-2 groups showed any clinical signs, whereas in the rLS1-XII-1 group some chickens became sick at day 3 post challenge. Viral shedding assessed in oral and cloacal swaps were in agreement with mortality and clinical signs. At the peak of viral shedding (4 dpc), positive swabs in rLS1 (1/14 for oral swabs and 3/14 cloacal swabs) and rLS1-XII-2 (0/14 and 1/14 for oral and cloacal swabs, respectively) groups were less than in rLS1-XII-1 group (9/13 and 9/13 for oral and cloacal swabs, respectively) (supplementary S1 Table). At 2nd and 4th dpc, 90% (9/10) and 100% (5/5) of the chickens in mock-vaccinated group were shedding virus (supplementary S1 Table).

**Fig 3.**
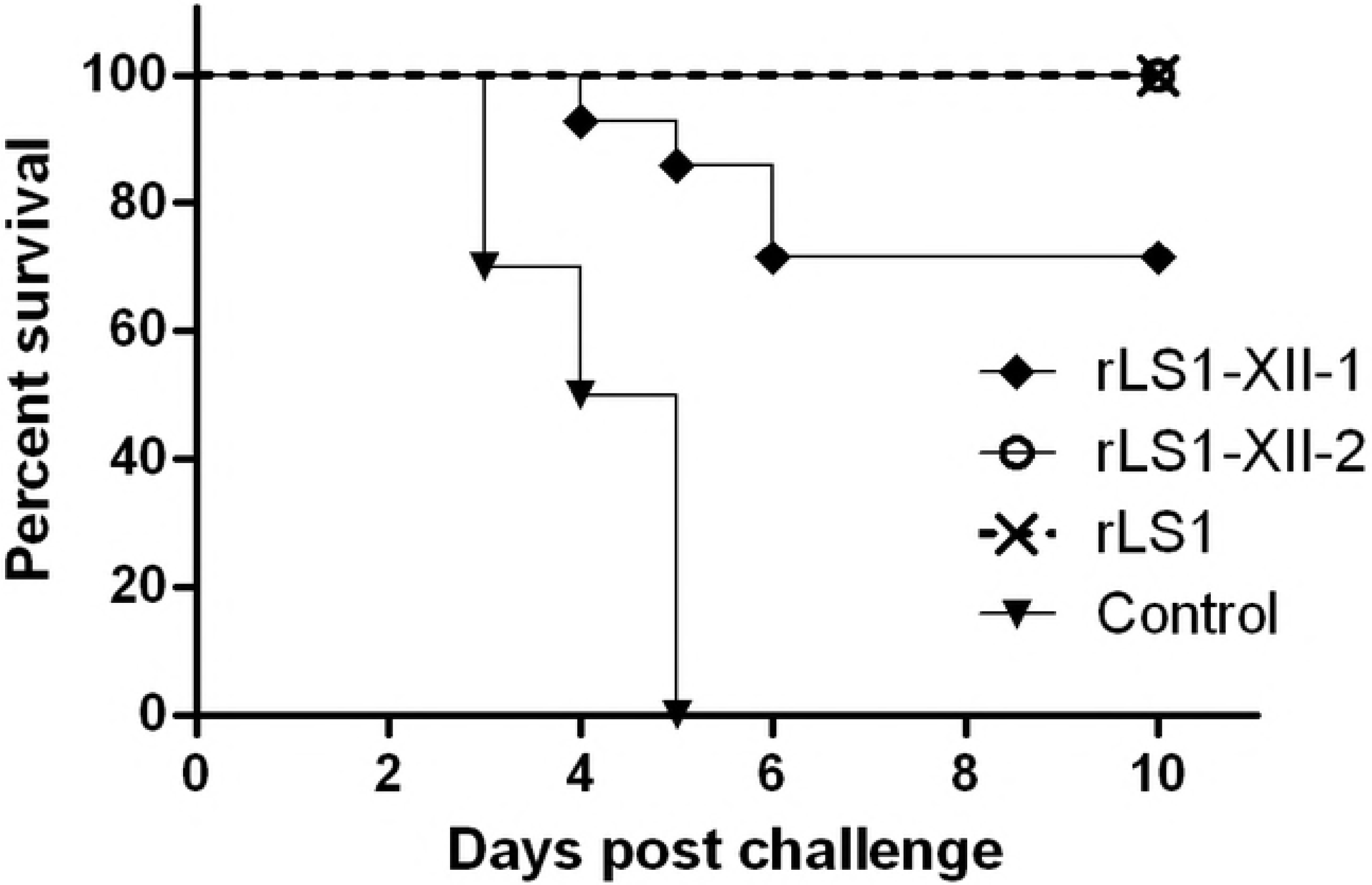
Survival rate after challenge with PP2011 (genotype XII) in chickens vaccinated with rNDVs. Chickens were immunized at 1 and 14 days of age. The challenge was performed at day 28 with 10^5^ LD_50_ per chicken via oculo-nasal route.

### Vaccination with rLS1-XII-2 was able to reduce viral shedding against a genotype XII challenge better than a commercial LaSota vaccine

In order to further determine whether there are differences between rLS1-XII-2 and a commercial LaSota vaccine strain, 7-day-old SPF chickens were immunized with only one dose of either of these viruses. Four weeks later, oculo-nasal challenge with PP2011 showed that both vaccine viruses protected 100% chickens against mortality, while PBS control group showed typical symptoms of velogenic NDV infection and had no survivors after day 5 post-challenge (Fig 4a). The shedding of challenged virus for immunized groups was assessed from oral and cloacal swabs at 2, 4 and 7 dpc. As shown in Table 3, the proportion of birds shedding challenge virus in mock-vaccinated group was the highest (between 80 and 100%). For LaSota vaccine group, 14-71 % of the swabs were positive, with a maximum peak at day 4 (7/14 for oral swabs and 10/14 for cloacal swabs). For rLS1-XII-2 vaccine group, only 0-29 % of oral and cloacal swabs were positive. In agreement with this, further quantitative analysis of positive swab samples showed that viral shedding in rLS1-XII-2 group tended to be significantly lower in oral and cloacal swabs compared to LaSota group (Fig 4b). Therefore, compared with the commercial LaSota vaccine, rLS1-is capable of reducing viral shedding after a challenge with a velogenic genotype XII strain.

**Table 3.** Frequency of isolation of challenge virus in vaccination groups.

| Group | Number of viral shedding chickens (positive/total) |  |  |  |  |  |
| --- | --- | --- | --- | --- | --- | --- |
|  | 2 dpc.<sup>(a)</sup> |  | 4 dpc |  | 7 dpc |  |
|  | Oral | Cloacal | Oral | Cloacal | Oral | Cloacal |
| rLS1-XII-2 | 0/14 | 2/14 | 2/14 | 4/14 | 0/14 | 2/14 |
| LaSota | 3/14 | 5/14 | 7/14 | 10/14 | 2/14 | 6/14 |
| PBS | 8/10 | 10/10 | 5/5 | 5/5 | NS<sup>(b)</sup> | NS |

(a) dpc = days post challenge
(b) NS = no survivors

**Fig 4.**
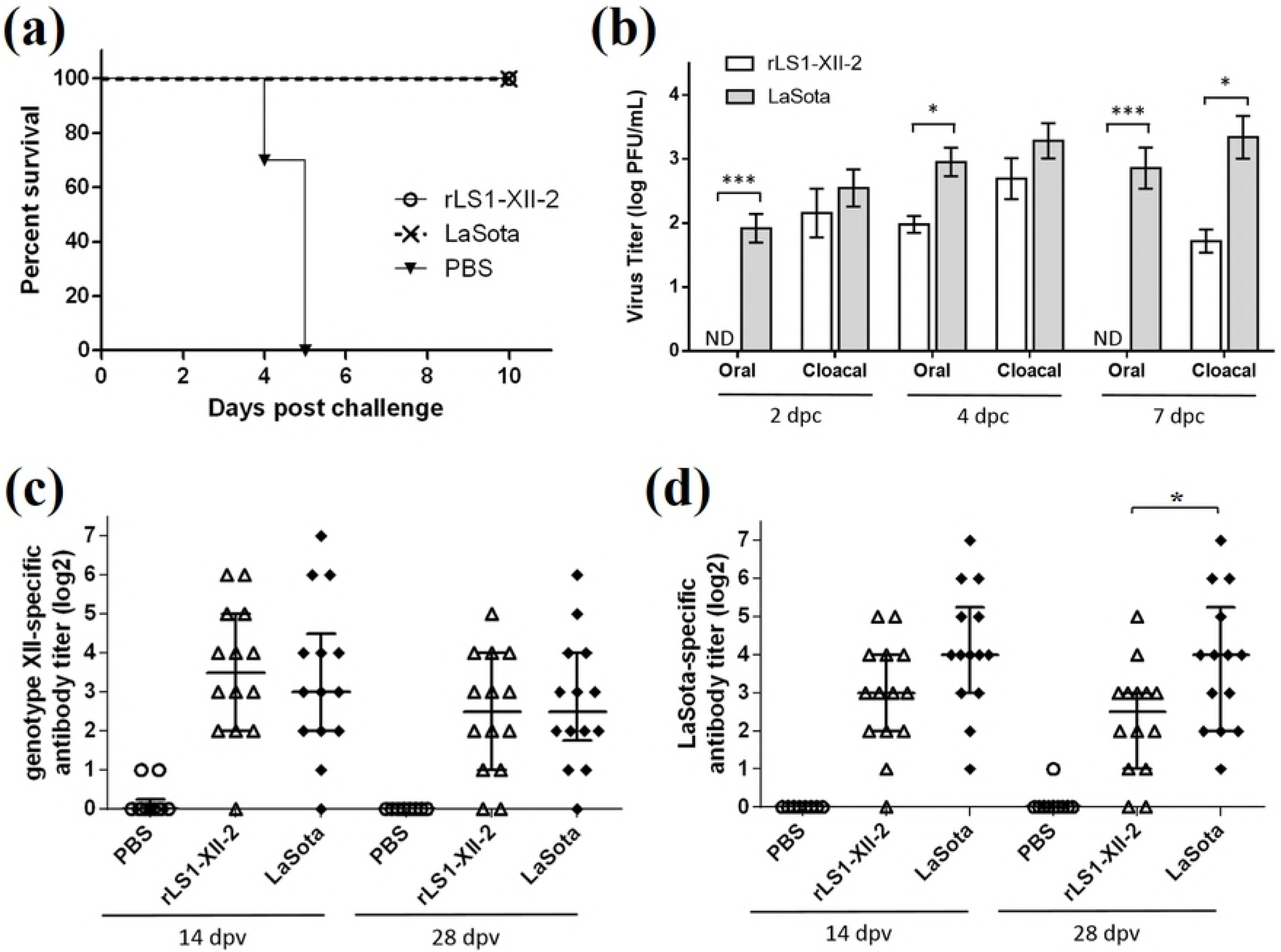
Comparison of the protection conferred by rLS1-XII-2 and LaSota strains against PP2011 (genotype XII) challenge. Chickens were immunized once at 7 days of age and the challenge performed 28 dpv with 10^5^ LD_50_/bird via oculo-nasal route. (a) Survival rate of vaccinated chickens showed no mortality in 10 days following the challenge. (b) Viral shedding from oral and cloacal swabs from chickens that were positive to challenge virus isolation. (c and d) Genotypes XII- and II-specific neutralizing antibodies in vaccinated chickens at 14 and 28 dpv. *p < 0.05 and ***p<0.001.

Genotype XII-specific serum neutralization titers taken at 14 or 28 dpv showed no difference between rLS1-XII-2 and LaSota strains (Fig 4c). However, the same samples tested for genotype II-specific neutralizing antibodies showed that LaSota group was significantly increased at 28 dpv (p = 0.031) and borderline significant at 14 dpv (p = 0.053) (Fig 4d).

## Discussion

Genotype XII is present in South America and is the prevalent circulating genotype in Peru [10,13,15]. In our country, current vaccination strategies have shown to be not sufficient to prevent outbreaks of the disease. This can be partially explained by antigenic differences between circulating and vaccine strains (see Table 1). Hence, improvements of vaccines and vaccination strategies are a priority. In this context, we decided to develop a genotype XII-matched, live attenuated vaccine candidate. First, we generated two chimeric NDVs, rLS1-XII-1 and -2. These viruses contain the F and HN proteins of the PP2011 strain (genotype XII) and the backbone of the parental rLS1 virus. For rLS1-XII-1, the complete ORFs of the F and HN proteins were replaced with those of the PP2011 strain, while, for rLS1-XII-2, only the ecto- and transmembrane domains of the F and HN proteins were replaced with those of the PP2011. Second, we tested the protection capacity of both viruses and the parental virus, obtaining that rLS1-XII-1 was not capable to fully protect SPF chickens against a genotype XII challenge. Finally, rLS1-XII-2 and a commercial LaSota vaccine were compared. Both strains fully protected SPF chickens, nonetheless, rLS1-XII-2 reduced significantly viral shedding. This places rLS1-XII-2 as a promising NDV vaccine candidate that would be suitable to better protect chickens against circulating genotype XII strains in Peru.

The presence of non-homologous cytoplasmic tails of F and HN proteins in genotype-matched NDV vaccine strains can produce an enormous impact on viral replication and, therefore, in protection. Genetically, the only difference between rLS1-XII-1 and -2 is the presence of homologous cytoplasmic tails in the latter. Nonetheless, rLS1-XII-1 has severely reduced the capacity of the virus to replicate (Fig 2). Other authors have developed genotype-matched NDV vaccines by replacing the complete ORFs of the F and HN protein genes with those of the strain of interest in LaSota backbone [21–24], as we did for rLS1-XII-1. In those studies, they found either greater [21,22] or similar [23,24] protection against homologous challenge compared to the parental recombinant LaSota. Although in those studies only genotypes VII and XI have been evaluated. Conversely, in this study the same strategy (rLS1-XII-1) resulted in a poorly replicating virus that was not able to prevent mortality, even with two vaccination doses, after a genotype XII challenge (see Fig 3). This was reverted when cytoplasmic tails of the F and HN proteins were restored to those original rLS1. The rLS1-XII-2 virus was able to prevent mortality as well as to reduce viral shedding better than a commercial LaSota vaccine (see Figs 4a and 4b and Table 3). Thus, one question arises, why previous studies have found at least the same (if not better) protection when complete F and HN ORFs were swapped with those of other genotypes? The most likely explanation is that F and/or HN cytoplasmic tails of the genotype XII strain PP2011 have a specific amino-acid(s) difference(s) that is not completely compatible with rLS1 backbone, affecting viral assembly or replication. Previously reported chimeric viruses do not have this specific amino-acid difference. The final consequence of this is a poorly replicating virus that cannot efficiently stimulate chicken immune system. This is reinforced by the fact that viruses with swapped cytoplasmic tails between NDV and an *Avian Avulavirus 2* cannot be recovered [30], or that disease severity can be negatively affected by the interaction between F and HN proteins with a heterologous M protein [31].

It has been shown that an efficient assembly and replication of the virus can be affected by removing key amino acids in the cytoplasmic tails of either the F or HN proteins [32,33]. Regarding the F protein, deletion of the last 6 amino-acids (positions 548 through 553), but not the last 4 amino-acids, of the cytoplasmic tail at the C-terminal region, impairs viral recovery [33], showing the importance of positions 548 and 549 for virus replication. Sequence analysis of the F protein of strains previously used to generate genotype-matched vaccines revealed that only PP2011 has a mutation in one of these positions with respect to LaSota (Fig 5a). This mutation (Ala549Thr) changes the charge of the amino acid from a hydrophobic to a polar amino-acid. Regarding the N-terminal cytoplasmic tail of the HN protein, positions 6 and 7 in the amino-acid sequence seem to be essential for viral replication [32]. The PP2011 strain has an amino-acid change in position 6 (Ser6Asp) which has a different nature to the one in rLS1, changing from an uncharged polar to a negative-charged amino-acid. While the other strains have different amino-acids at this position (Ser6Asn or Ser6Thr), these do not change the polar nature of this amino acid position (see Fig 5b). Gln7Lys mutation in position 7 is also unique for PP2011, although the nature of this change (from uncharged polar to basic) is the same to the ones present in the other analyzed strains (Gln7Arg). Despite that, the above described amino-acid changes are the most likely to cause the phenotype discrepancies between rLS1-XII-1 and -2, it is unclear which mutation(s) is responsible for the observed phenotype. Besides, we cannot rule out the possibility that the other mutation(s) in the cytoplasmic tails of either F or HN proteins may play a role in virus fitness. Further studies should determine the individual effect of each of these mutations and reveal key amino-acid positions for viral assembly and replication.

**Fig 5.**
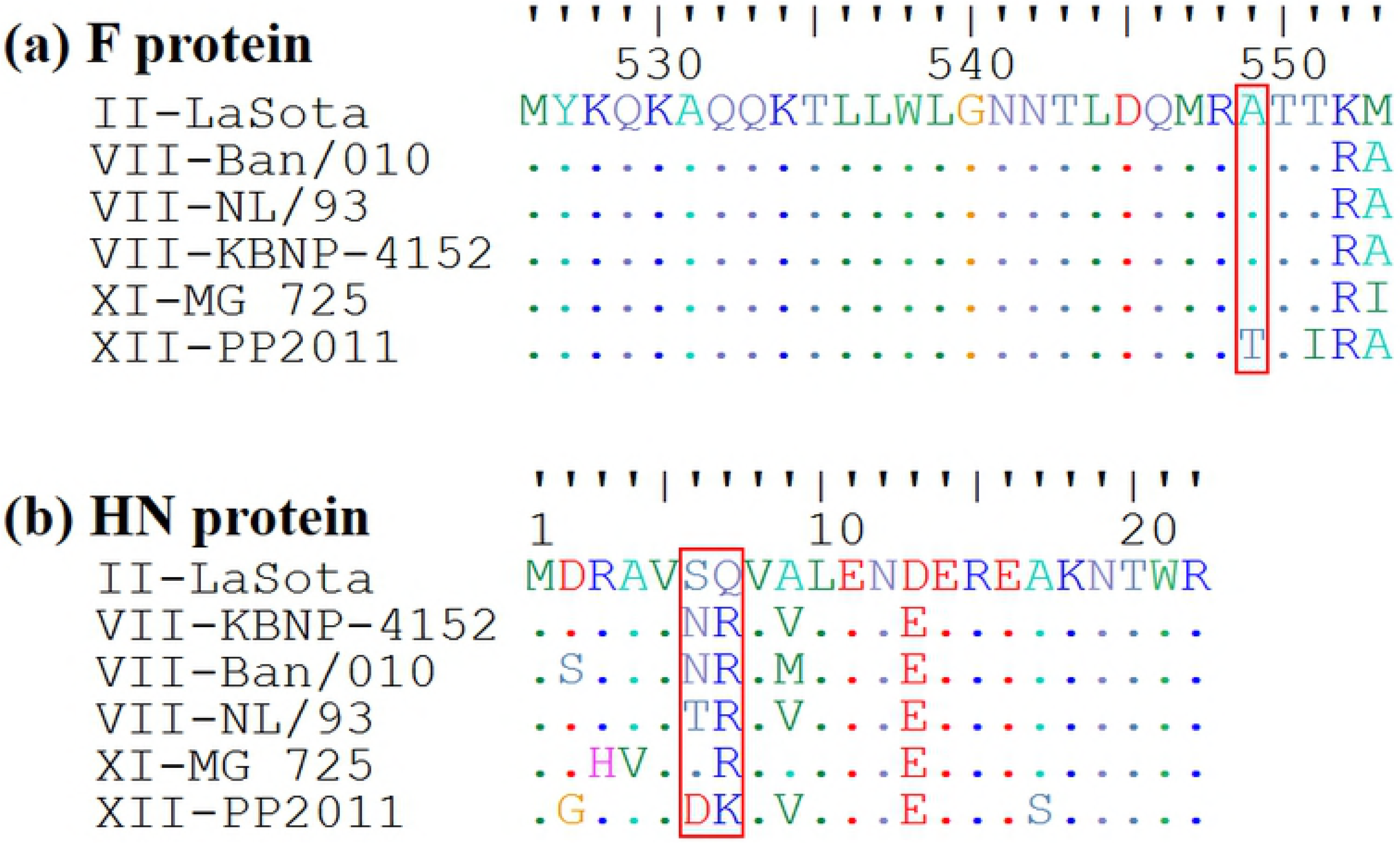
Amino-acid sequence alignment of F (a) and HN (b) cytoplasmic tails from NDVs used to generate chimeric viruses. F and HN sequences from strains Ban/010 (Acc. N° HQ697254) [22], NL/93 (Acc. N° JN986837) [23], KBNP-4152 (Acc. N° DQ839397) [21] and MG_725 (Acc. N° HQ266602) [24] were previously used to generate chimeric NDVs in LaSota (Acc. N° AF077761) backbone. PP2011 (Acc. N° KR732614) [14] F and HN cytoplasmic tail sequences were used in this study to generate rLS1-XII-1 strain in the rLS1 backbone. Amino-acid sequences of F and HN cytoplasmic tails are exactly the same for LaSota and rLS1.

Increased serum neutralizing antibodies are not a requirement for reduction in viral shedding. In our second experiment, we showed that compared with LaSota, rLS1-XII-2 vaccine reduced viral shedding in both the number of shedding birds and the quantity of secreted virus per positive chicken (see Table 2 and Fig 4b). Nevertheless, we did not find increased genotype XII-specific neutralizing antibody titers in rLS1-XII-2 (Fig 4c). These results seem contradictory compared with previous experiments [18,19,22,25,27], but to explain this apparent discrepancy, we suggest two possibilities: 1) Local immunity was increased in respiratory and gut tracts in chickens vaccinated with rLS1-XII-2 compared to those vaccinated with LaSota. Given that rLS1-XII-2 has the surface proteins of the genotype XII, it could have a tropism towards respiratory and gut tracts [15], similar to the challenge strain, while LaSota replicates almost exclusively in the respiratory tract [34]. This can elicit a greater local and more specific response, evidenced by, for example, the increased number of genotype XII-specific resident memory T and B cells or the elevated levels of specific-IgA in mucosal tissues, but not in serum. 2) Cellular immune response may help to clear the infection rapidly, avoiding or reducing viral shedding. Chickens vaccinated with rLS1-XII-2 are exposed to more similar epitopes of the PP2011 challenge strain than chickens vaccinated with LaSota vaccine. These epitopes can be presented to CD4+ or CD8+ T cells via major histocompatibility complex class-I (MHC-I) and class-II (MHC-II) and start to clear the infection. Nonetheless, neither of these mechanisms were assessed in this study and we cannot rule out other possible explanations.

In conclusion, we have developed a recombinant NDV vaccine candidate capable of fully protecting chickens and reduce viral shedding against a genotype XII homologous challenge. Furthermore, we have showed the importance of cytoplasmic tails in replication to generate competent vaccine viruses. Therefore, the balance between replication and antigenicity should be taken into account for developing efficient genotype-matched vaccines.

## Acknowledgements

The authors would like to thank Edison Huaccachi, James Quispe and Julia Garayar for their excellent technical assistance.

## Author’s contributions

RI-L, AC, MF-D and VNV contributed to conception and designed the research. RI-L, AC and KC acquired the data of the study. RI-L analyzed the data, and all authors interpreted them. RI-L and VNV drafted the paper, and all authors revised it critically, read and approved the final manuscript.

## Supporting information

S1 Table. Frequency of isolation of challenge virus in vaccinated groups of the first experiment

